# Innovative Use of Depth Data to Estimate Energy Intake and Expenditure in Adélie Penguins

**DOI:** 10.1101/2024.07.02.601650

**Authors:** Benjamin Dupuis, Akiko Kato, Olivia Hicks, Danuta M. Wisniewska, Coline Marciau, Frederic Angelier, Yan Ropert-Coudert, Marianna Chimienti

## Abstract

Energy governs species’ life histories and pace of living, requiring individuals to make trade-offs. However, measuring energetic parameters in the wild is challenging, often resulting in data collected from heterogeneous sources. This complicates comprehensive analysis and hampers transferability within and across case studies. We present a novel framework, combining information obtained from eco-physiology and bio-logging techniques, to estimate both energy expended and acquired on 48 Adélie penguins (*Pygoscelis adeliae*) during the chick-rearing stage.

We employ the machine learning algorithm random forest (RF) to predict accelerometry-estimated foraging behaviour using depth data (our proxy for energy acquisition). We also build a time-activity model calibrated with doubly labelled water data to estimate energy expenditure.

Using depth-derived time spent diving and amount of vertical movement in the sub-surface phase, we accurately predict energy expenditure (R² = 0.70). Movement metrics derived from depth data modelled with the RF algorithm were able to accurately (accuracy = 0.82) detect the same foraging behaviour predicted from accelerometry. The RF more accurately predicted accelerometry-estimated time spent foraging (R² = 0.81) compared to historical proxies like number of undulations (R² = 0.51) or dive bottom duration (R² = 0.31).

The proposed framework is accurate, reliable and simple to implement, enabling to couple energy intake and expenditure, which is crucial to further assess individual trade-offs. We provide universal guidelines for predicting these parameters based on widely used bio-logging technology in marine species. Our work allows us to revisit historical data, to study how long-term environmental changes affect animals’ energetics.

**Summary statement:** Using machine learning, we estimated energy expenditure and foraging activity of free-ranging Adélie penguins using depth data recorded with bio-logging devices.

## 1. Introduction

Energy is the fundamental currency shaping animals’ life-history strategies (Burger et al., 2019; Kressler et al., 2023; Pontzer and McGrosky, 2022). Individuals within populations acquire and spend energy to fuel activities such as movement, body maintenance, thermoregulation and reproduction (Gower et al., 2008; Noakes et al., 2013; Steinhart et al., 2005). As energy acquisition from the environment is limited, individuals’ performances and trade-offs in energy allocation directly impacts life-histories traits like survival and reproduction, and therefore fitness and population processes (Brown et al., 2004; Mogensen and Post, 2012; Morano et al., 2013). In addition to fuelling all physiological processes, energy acquisition (i.e. foraging) itself requires physical activity and therefore energy expenditure (Pontzer and McGrosky, 2022). Environmental variations in biotic (e.g., in prey availability) and abiotic factors, and animal internal states can affect how energy is spent and acquired, and lead to adjustments in foraging strategies (Byrne et al., 2022; Chevallay et al., 2022; Egert-Berg et al., 2021).

In order to understand individual trade-offs, it is important to consider both energy acquisition and expenditure within the same framework. Yet, simultaneously measuring energy intake and expenditure in wild animals is challenging. Numerous species forage in areas almost impossible to sample (*i.e.* deep oceans, remote land, sky), making observations difficult. Aapproaches in eco-physiology, like doubly-labelled water (DLW), respirometry, heart-rate monitors, or oesophagus temperature sensors have proven useful in providing data to validate energy expenditure and intake (Froget et al., 2004; Hicks et al., 2020; Nagy et al., 1999; Ropert-Coudert et al., 2001). However, these approaches are costly, logistically difficult, invasive and, therefore, challenging to consistently use in long-term species monitoring programs. Given the difficulties in simultaneously measuring both energy expenditure and energy acquisition in wild animals, information comes from different data types (English et al., 2024) making long-term analysis or study comparisons difficult.

Recent advances in bio-logging technologies allowed researchers to validate the use of accelerometers to accurately estimate animals’ energetics across a wide range of taxa and habitats (English et al., 2024; Wilson et al., 2020). Top predators integrate information from the bottom to the top of the food webs. As such, they are recognized as sentinels of global change, and allow understanding the impact of environmental changes on ecosystems (Hazen et al., 2019; Sergio et al., 2008). Marine top predators are of special interest because they live in environments where global changes have rapid and important consequences on ecosystems (Sydeman et al., 2015). They are often both wide-ranging and conspicuous (compared to lower-marine trophic levels) (Hazen et al., 2019). By recording tri-axial body acceleration at high resolution (25-100 Hz) on top predator species such as white sharks (Watanabe et al., 2019), southern elephant seals (Gallon et al., 2013) or penguins (Kokubun et al., 2011), it is possible to detect prey capture attempts and to classify behaviours. Such methodology also allows us to estimate energy expenditure across foraging trips when validated with the DLW technique for example (Chimienti et al., 2016; Del Caño et al., 2021; Hicks et al., 2020). Yet, because of their novelty, historical time series of accelerometer data are often not available.

On the contrary, time-depth recorders (TDRs) have been extensively used on diving predators since the 90’s (Ropert-Coudert et al., 2009) to reconstruct dive profiles, investigate foraging behaviour, and estimate energy expenditure of diving marine predators (Chappell et al., 1993a, 1993b; Viviant et al., 2014). Despite lower accuracy, data collection via TDRs presents several advantages compared to data collected with accelerometer tags. For instance, TDRs are more suitable for extended recording periods (months or years of continuous recording). TDRs record data at lower resolution (usually 1Hz or coarser) requiring reduced memory space and battery consumption, and do not need to be placed at or near the centre of the mass of the animal like accelerometers. TDRs can be used to study the foraging behaviour of diving marine predators over several consecutive annual cycles, such as in Adélie penguins (Lescroël et al., 2023). Moreover, the coarser data resolution, compared to accelerometer data, generates smaller datasets which require less analytical and computational power. These advantages are very important when working on long-term species monitoring programmes and when aiming to link individual behaviour to fitness (reproduction and survival), and ultimately, population dynamics.

Several TDR-derived time-activity budget models or calibrations of metabolic rate with DLW measurement have been built across marine species to estimate energy expenditure (Chappell et al., 1993b; Chivers et al., 2012). Dive metrics such as bottom phase duration and number of undulations within dives were used as proxies of foraging activity (Bost et al., 2007; Lescroël et al., 2021; Viviant et al., 2014). Yet, validation of such metrics is rare, and recent papers tend to show that these metrics alone do not effectively reflect the foraging activity of marine predators (Allegue et al., 2023; Brisson-Curadeau et al., 2021).

We examined this question in Adélie penguins, one of the most abundant Antarctic seabird species and ecosystem sentinel of the Southern Ocean (Barbraud et al., 2020; Forcada and Trathan, 2009). This species mostly forages on two species of krill (*Euphausia superba* and *E. crystallorophias*) (Ratcliffe and Trathan, 2012), and are therefore highly dependent on sea-ice conditions (Kokubun et al., 2021; Michelot et al., 2020). Bottom phase duration and undulations have been extensively used to describe its foraging activity. Yet, the thresholds used to define these two parameters are often different across studies. Bottom phase is sometimes considered spanning from the first to last time vertical velocity was < 0.25 m.s^-1^ (Ropert-Coudert et al., 2007), sometimes it is considered to be below the 40% deepest part of a dive (Lescroël et al., 2021), and it sometimes spans from the first to last undulation (Bost et al., 2007). Similarly, calculations of the number of undulations within a dive are derived either from changes in vertical velocity alone or with the latter in addition to different intensity thresholds (Bost et al., 2007; Lescroël et al., 2021; Ropert-Coudert et al., 2001). Creating a simple, validated and objective framework to study its foraging behaviour based on depth data could therefore ease study comparisons.

The colony located on île des Pétrels, Antarctica, has been extensively monitored since the 1960’s. Since the mid 1990’s, TDRs were also regularly deployed, followed by accelerometers since 2016. Hence, only eight years of accelerometry data are currently available, compared to twenty-five for TDR. During the 2018-19 breeding season, breeding Adélie penguins were fitted with loggers recording both accelerometry and TDR data (Hicks et al., 2020). Importantly, DLW measurements were collected from these individuals, allowing the calculation of accurate data on energy expenditures. We take advantage of this diverse ecological data collection to develop a framework allowing estimation of energy balance of Adélie penguins from depth data. We use DLW measurements and TDR data to predict energy expenditure from depth data only. We combine behavioural classification based on accelerometry (Chimienti et al., 2022) and the power of machine learning (Pichler and Hartig, 2023) to estimate foraging activity on solely depth data. We compare our results with other methods classic TDR metrics and accelerometers to answer the following research questions: 1. Can depth data be used to predict energy expenditure of marine predators? and 2. Can machine learning help estimate foraging activity of marine predators from depth data without relying on arbitrary thresholds? 3. Furthermore, since foraging is a costly behaviour, we test whether our framework can reproduce the pattern of DEE increase with time spent foraging.

## 2. Material and methods

### 2.1. Data collection

The study colony is located on Ile des Pétrels, next to the Dumont D’Urville research station, in Adélie Land (66°40’ S; 140°01’ E). From the 21^st^ December 2018 to the 11 January 2019, 58 breeding Adélie penguins (24 females and 34 males) were tracked and monitored. All individuals were in their chick guarding stage, where parents alternate mostly 1-day trips at sea to forage and feed their chicks (Ainley, 2002). Individuals were captured at their nest when both parents were present. To limit disturbance, we only captured one of the partners for each nest. We performed molecular sexing at CEBC as previously described (Marciau et al., 2023) to confirm the sex of each individual *a posteriori*.

This study was approved and authorized by the ethics committee number 084 of the Terres Australes et Antarctiques Françaises (TAAF), Comité d’Environnement Polaire and Conseil National de la Protection de la Nature. All experiments were performed in accordance with the guidelines of these committees.

### 2.2. Logger deployment & Data preparation

Data loggers (Axy-Trek, Technosmart, Italy, 40 x 20 x 8 mm, 14g, less than 0.5 % of individuals mass) recording tri-axial acceleration at 100 Hz and pressure at 1 Hz were deployed on the central back region of breeding Adélie penguins and secured using waterproof adhesive Tesa tape and two Colson plastic cable ties. Deployment duration ranged from 46.53 hours to 78.43 hours with an average of 54.82±1.43 hours. Trip duration (from first to last dive) ranged from 12.24 hours to 31.94 hours with an average of 22.60±0.98 hours.

Upon recovery, data were downloaded and processed using the R programming language. After calculating depth (± 0.1 m) from pressure (Leroy and Parthiot, 1998), a custom function (see available code) was used to perform the zero-offset correction. Static acceleration was calculated by smoothing each axis over a 1-s period. Then, Dynamic Body Acceleration (DBA) was calculated by subtracting static acceleration from raw acceleration value. Vectorial DBA (VeDBA) was calculated as the square root of the sum of the squares DBA of the three axes.

Daily energy expenditure (DEE, kJ/day) was measured using the doubly labelled water technique, as described in Hicks et al. (Hicks et al., 2020). DEE was corrected by the mass of the individual upon equipment to get a mass specific DEE (kJ/g/day).

### 2.3. Behavioural assignment on depth data for energy expenditure estimation

For each individual, we calculated the time spent in a given behaviour across logger deployment. Each behaviour was defined based on depth data. An individual was considered diving when it was below 2m, swimming/resting on the water when it was above 1m, and in subsurface or proposing when it was between 1 and 2m (fig. 1). Penguins were considered on land between logger deployment and first dive, last dive and logger recovery, and when they were on the surface for more than 6 hours. In addition to duration, we quantified the overall sum of vertical movement for each phase (except land).

**Figure 1.**
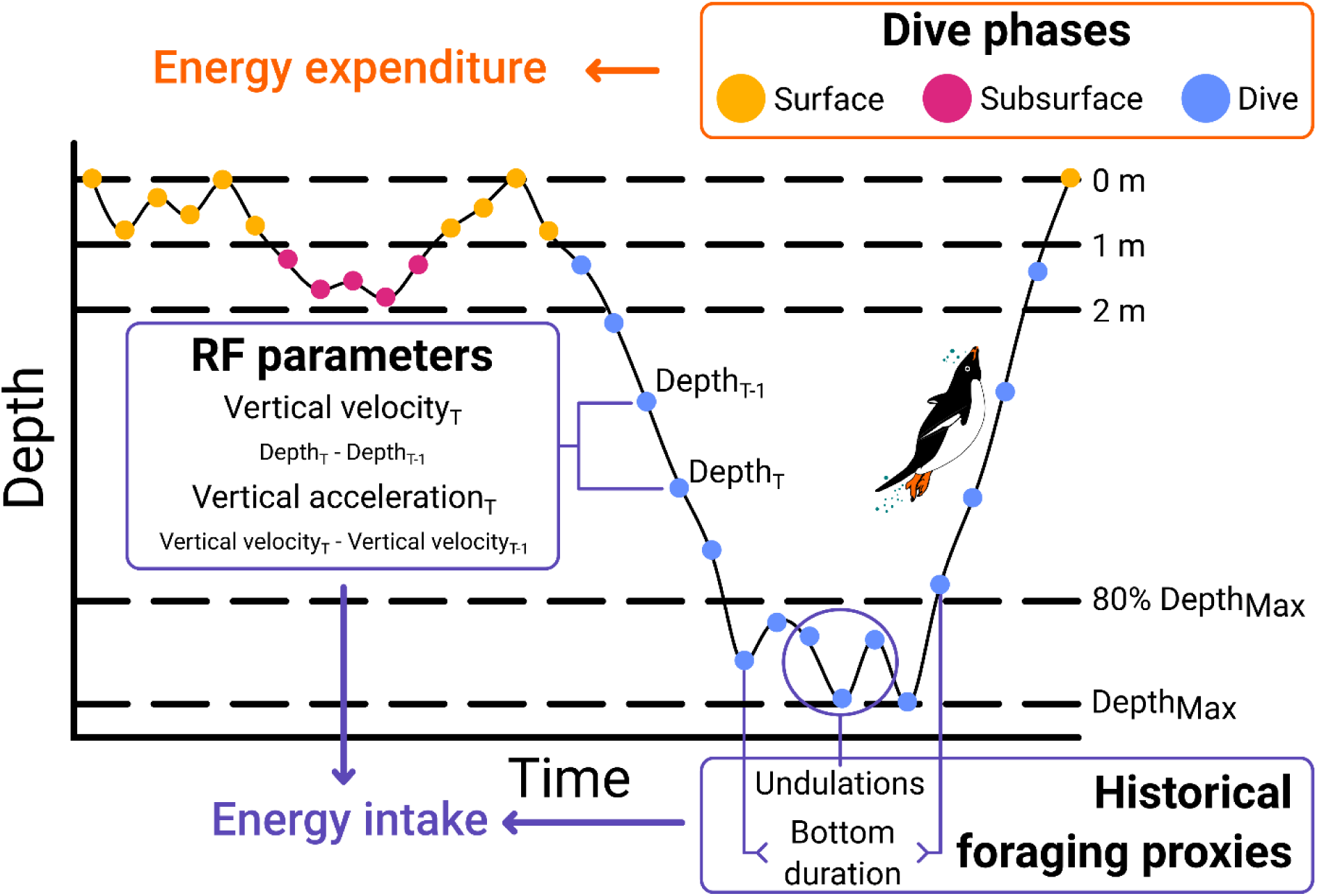
Conceptual visualization of the depth-derived parameters used to estimate energy balance of Adélie penguins.

### 2.4. Machine learning approach for foraging detection

In addition to raw depth value, we derived a set of parameters from depth data to feed our machine learning algorithm and describe foraging behaviour. For each dive, we calculated the maximum depth and the dive duration (continuous period below 2m). To estimate fine-scale movement, we calculated the vertical velocity as a derivative of depth between time T and T_-1_ (eqn. 1) and the vertical acceleration as the derivative of velocity between time T and T_-1_ (eqn. 2, fig. 1) for each data point (1Hz resolution). To estimate broader-scale movement, we calculated the rolling mean and standard deviation of vertical velocity and vertical acceleration over a 5-seconds period.

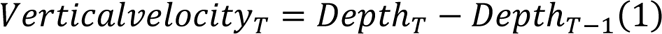

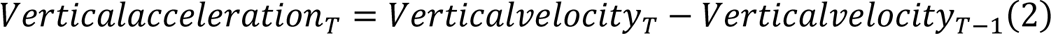

Using a random forest algorithm (RF), we tried to identify periods of foraging in the diving behaviour of Adélie penguins. As a reference, we used the accelerometry-based behavioural classification from Chimienti *et al*. (Chimienti et al., 2022). As this classification was done at 25 Hz, we summarised it at 1 Hz. To be conservative, we considered 1 Hz data point as foraging when at least half of the corresponding 25 Hz were labelled as foraging (Machado-Gaye et al., 2024), regardless of the proportion. Before running the RF, we performed variable selection to reduce the size of our model. We filtered variables based on correlation factor and variables importance measure (VIM, from “Boruta” R package (Kursa and Rudnicki, 2010)). Whenever two variables were highly correlated (>0.8), the one with the lowest VIM was removed from the RF.

Remaining variables were used in a RF built using “tidymodels” (Kuhn and Wickham, 2020) and “ranger” (Wright and Ziegler, 2017) R packages. We randomly selected and assigned half of the deployments to train the RF, while the other half was kept to test model performance. Train and test dataset had the same sex-ratio. We trained the RF over 6 variables and parametrised it on different numbers of trees (*i.e.* 50, 100, 500, 1000). We also tuned the *mtry* parameter, indicating the number of variables randomly sampled as candidates at each split, between 1 and 6 for each number of trees. Finally, the training dataset was further split into training (75%) and testing (25%) and a five-fold cross validation procedure performed on the final training data. The best model was selected based on two widely used metrics, namely accuracy and Area Under the Receiver Operating Characteristic (ROC) Curve (AUC).

### 2.5. Statistical analysis of energetics

All statistical analyses and data manipulations were performed in R version 4.3.1 (R Core Team, 2023). Unless stated otherwise, means ± SE are provided. All model assumptions were checked using the plot_model() function for the “sjplot” (Lüdecke, 2023) R package.

To estimate energy expenditure from depth data, we modelled the DLW-derived DEE using linear models with sex, time spent in each behaviour, and sum of vertical movement over each behaviour at the daily level. We compared all combinations using Akaike’s information criterion corrected for small sample size (AICc) and Bayesian information criterion (BIC). To estimate the prediction power of our model, we implemented a bootstrap procedure to limit the effect of our reduced sample size. Over 1000 iterations, the dataset was randomly separated in a train (50%) and test (50%) dataset. After fitting the best model to the train dataset, we compared its depth-derived DEE to the DLW-derived on the test dataset. For each coefficient, we calculated a 95% confidence interval to estimate model stability.

To estimate foraging activity from depth data, we modelled at the daily level the accelerometry-derived time spent foraging at-sea T_foraging_ with the RF-derived T_foraging_, the RF-derived number of prey catching attempts (PCA, *i.e.* continuous foraging period), and two widely-used proxies to quantify foraging intensity, namely bottom duration (*i.e.* time spent in the 20% deepest part of a dive (Bestley et al., 2015; Carter et al., 2017) and number of undulations (*i.e.* change from a negative to positive or positive to negative vertical velocity in the bottom phase (Lescroël et al., 2021). Again, all model combinations were compared using AICc and BIC.

## 3. Results

### 3.1. Energy expenditure estimation

The most parsimonious model was able to reliably predict DLW-estimated DEE (adjusted R² = 0.70, fig. S1). This model retained only 3 parameters: time spent diving, amount of vertical movement in the sub-surface phase, and sex (eqn. 3, table 1). All models within a ΔAICc of 2 contained both time and movement-based parameters (table 1). In all models, even when it was retained, sex was never significant. In comparison, the most parsimonious model performed significantly better than the best time-activity model (ΔAICc = 3.28, table S1) and the null VEBDA model (ΔAICc = 8.15).

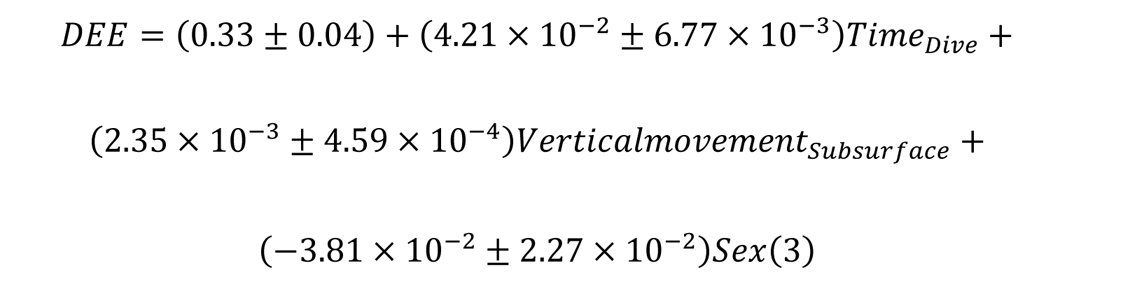

**Table 1.** Model selection for DLW-derived DEE model. Only models within ΔAICc of 2 are displayed, as well as the best time-activity model (*), and a null model (**).

| Formula | Rank | AICc | BIC | $\Delta AICc$ | Weight | Adj. R<sup>2</sup> |
| --- | --- | --- | --- | --- | --- | --- |
| DEE ~ T<sub>Dive</sub> + Vertical | 4.00 | - | - | 0.00 | 0.41 | 0.70 |
| Movement<sub>Sub-surface</sub> + Sex |  | 100.0 | 92.6 |  |  |  |
|  |  | 0 | 8 |  |  |  |
| DEE ~ T<sub>Dive</sub> + Vertical | 3.00 | - | - | 0.42 | 0.33 | 0.68 |
| Movement<sub>Sub-surface</sub> |  | 99.58 | 93.4 |  |  |  |
|  |  |  | 7 |  |  |  |
| DEE ~ T<sub>Dive</sub> + T<sub>Sub-surface</sub> + | 5.00 | - | - | 1.64 | 0.18 | 0.70 |
| Vertical Movement<sub>Sub-</sub> |  | 98.36 | 89.9 |  |  |  |
| surface + Sex |  |  | 3 |  |  |  |
| * DEE ~ T<sub>Dive</sub> + T<sub>Sub-surface</sub> | 3.00 | - | - | 3.28 | 0.08 | 0.66 |
|  |  | 96.72 | 90.6 |  |  |  |
|  |  |  | 1 |  |  |  |
| ** DEE ~ Mean<sub>VeDBA</sub> | 2.00 | - | - | 8.15 | 0.01 | 0.61 |
|  |  | 91.85 | 87.1 |  |  |  |
|  |  |  | 0 |  |  |  |

The bootstrap procedure confirmed the robustness of our model to predict DEE from depth data (fig. 2, ρ = 0.81 [0.64; 0.91], R² = 0.69 [0.45; 0.85]) and was coherent with model coefficient calculated on the full dataset.

**Figure 2.**
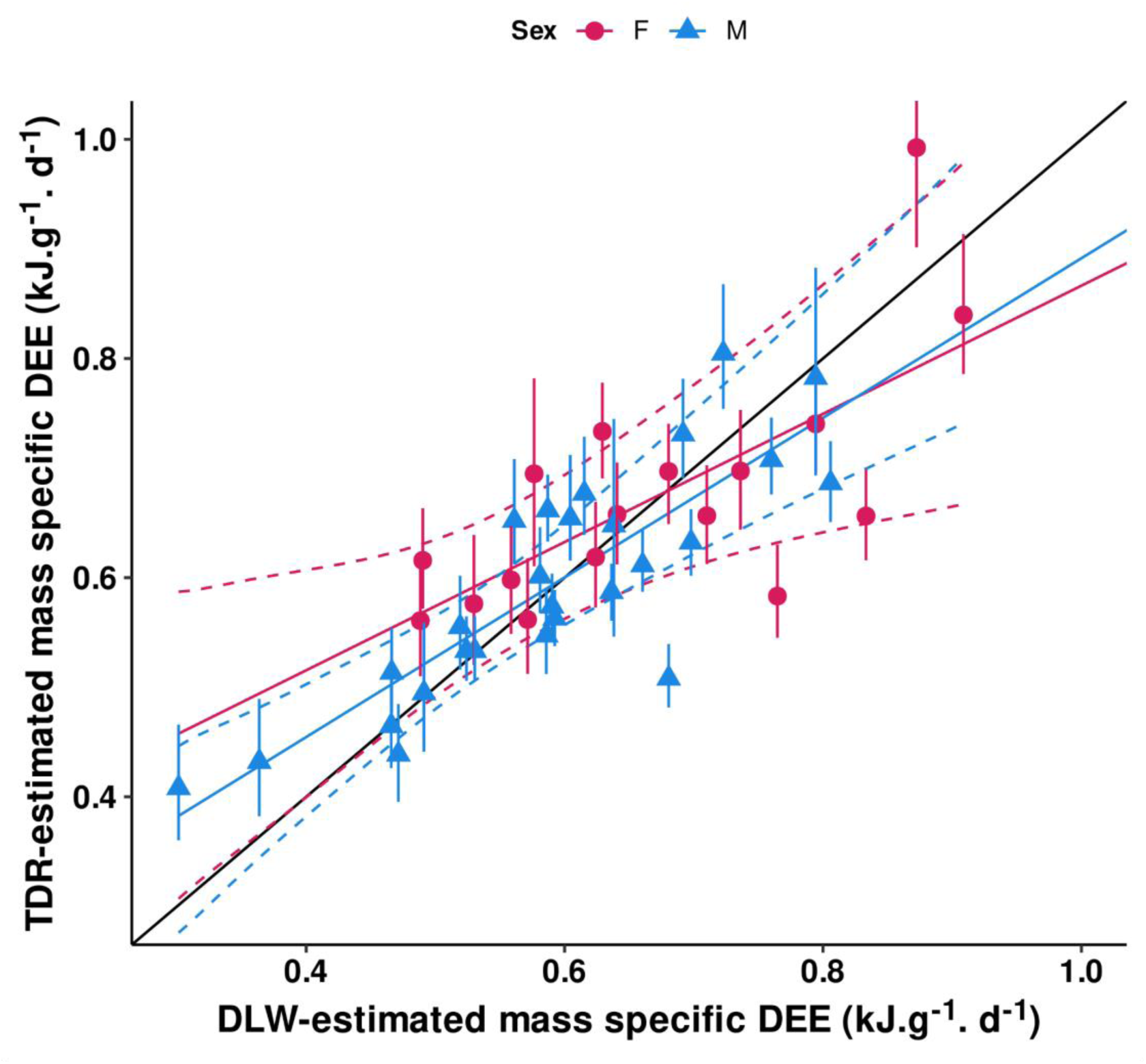
Relation between TDR and DLW-estimated DEE. Dashed lines and error bars represent 95% confidence intervals. The black line represents the y = x slope.

### 3.2. Random forest

During model tuning, accuracy and AUC plateaued when *mtry* reached 3 and *ntrees* 1000. With these parameters, the Out-Of-Bar (OOB) error estimated the RF algorithm was 0.03, indicating the good predictive power of our model. In decreasing order of importance, retained variables were the rolling SD of vertical acceleration, the rolling mean of vertical velocity, the depth, the dive duration, the rolling mean of verticalacceleration, and vertical acceleration (fig. S2). Overall accuracy of the model was high, with a balanced accuracy of 0.83, specificity of 0.95 and sensitivity of 0.72. When evaluating the fine scale detection of foraging events, we observed that the model tends to group short consecutive PCA. Nonetheless, the overall foraging duration was very similar to the accelerometry-based reference label and our RF forest algorithm detected foraging periods during the ascend phase of the dives (fig. 3).

**Figure 3.**
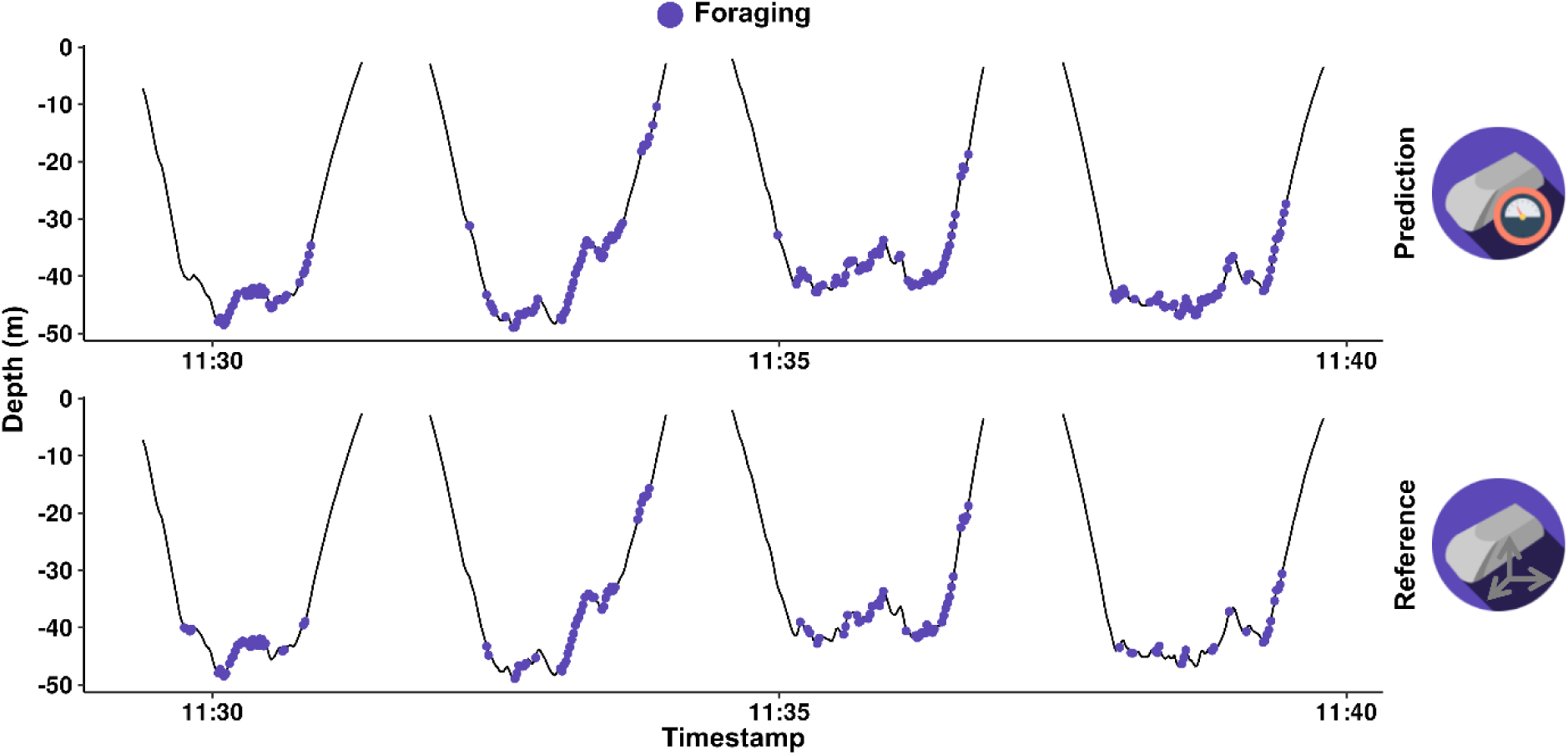
Dive profile of Adélie penguin. A. With the RF algorithm based on depth data. B. With the reference accelerometry-based classification. Purple dots represent foraging.

### 3.3. Foraging activity estimation and comparison with other proxies

We investigated if our RF algorithm could predict time spent foraging at-sea and compared it with historical proxies. Our RF efficiently predicted accelerometry-based time spent foraging over a foraging trip from TDR-data (R² = 0.81, AICc = 18.22). This model was significantly better at predicting the reference accelerometry-based time spent foraging than bottom phase duration (R² = 0.51, AICc = 40.76) and number of undulations (R² = 0.32, AICc = 48.78 table 2). Using multiple foraging proxies did not significantly increase the predictive power of our model (fig. 4, table S2). All proxies were predicting time spent foraging better than the null model with trip duration only (R² = 0.04, AICc = 57.15)

**Figure 4.**
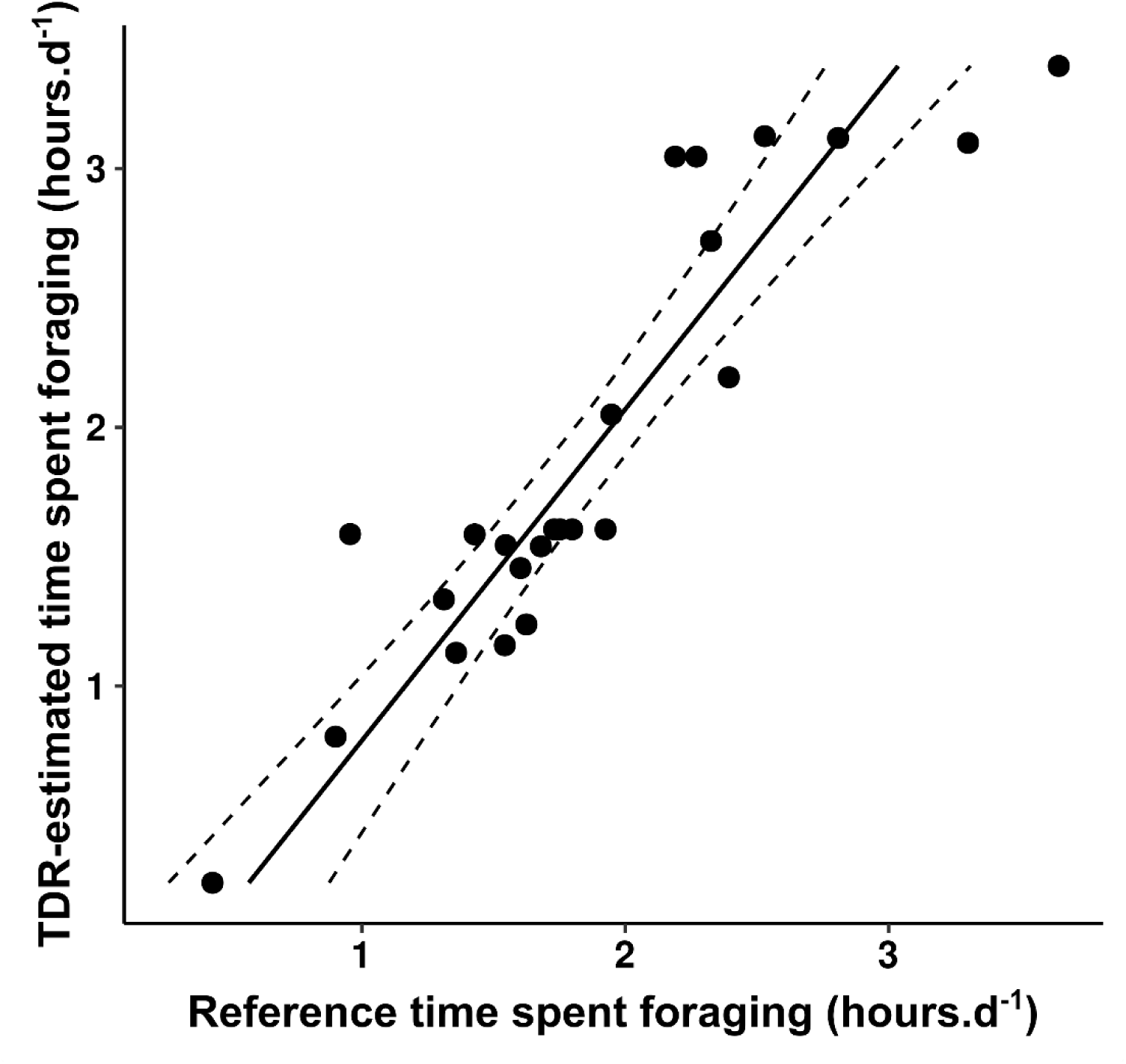
Output from the most parsimonious model to predict time spent foraging per day at sea. Reference time spent foraging was calculated from high-resolution tri-axial accelerometry data. Dashed lines represent the 95% confidence interval around the regression.

**Table 2.** Model selection for predicting Accelerometry based time spent foraging. Only models within ΔAICc of 2 are displayed, as well as three null models, with only bottom phase duration (*), number of undulations (**), and trip duration (***).

| Formula | Ran<br>k | AIC<br>c | BIC | $\Delta AICc$ | weig<br>ht | adj.r.sqrd |
| --- | --- | --- | --- | --- | --- | --- |
| RefTime spent foraging ~ | 2.00 | 18.2 | 20.5 | 0.00 | 0.34 | 0.81 |
| PredTime spent foraging |  | 2 | 6 |  |  |  |
| RefTime spent foraging ~ | 3.00 | 18.3 | 20.9 | 0.09 | 0.33 | 0.82 |
| PredTime spent foraging +<br>TimeBottom phase |  | 1 | 1 |  |  |  |
| * RefTime spent foraging ~<br>TimeBottom phase | 2.00 | 40.7 | 43.1 | 22.54 | 0.00 | 0.51 |
|  |  | 6 | 0 |  |  |  |
| ** RefTime spent foraging ~<br>NUndulations | 2.00 | 48.7 | 51.1 | 30.56 | 0.00 | 0.32 |
|  |  | 8 | 2 |  |  |  |
| *** RefTime spent foraging ~<br>Trip duration | 2.00 | 57.1 | 59.4 | 38.93 | 0.00 | 0.04 |
|  |  | 5 | 8 |  |  |  |

### 3.4. At-sea energetics of Adélie penguins

On average, penguins spent 1.84 ± 0.11 hours foraging per trip (0.81 ± 0.05 hours per day). Over that same period, individuals expend on average 0.63 ± 0.02 kJ.g^-1^ per day, which corresponds to a mean DEE of 2983.01 ± 72.99 kJ. Penguins spending more time foraging per day displayed comparatively higher DEE (fig. 5, table S3, estimate = 0.11 ± 0.04, p < 0.01). For a given foraging duration, females had higher DEE than males (estimate = - 0.09 ± 0.03).

**Figure 5.**
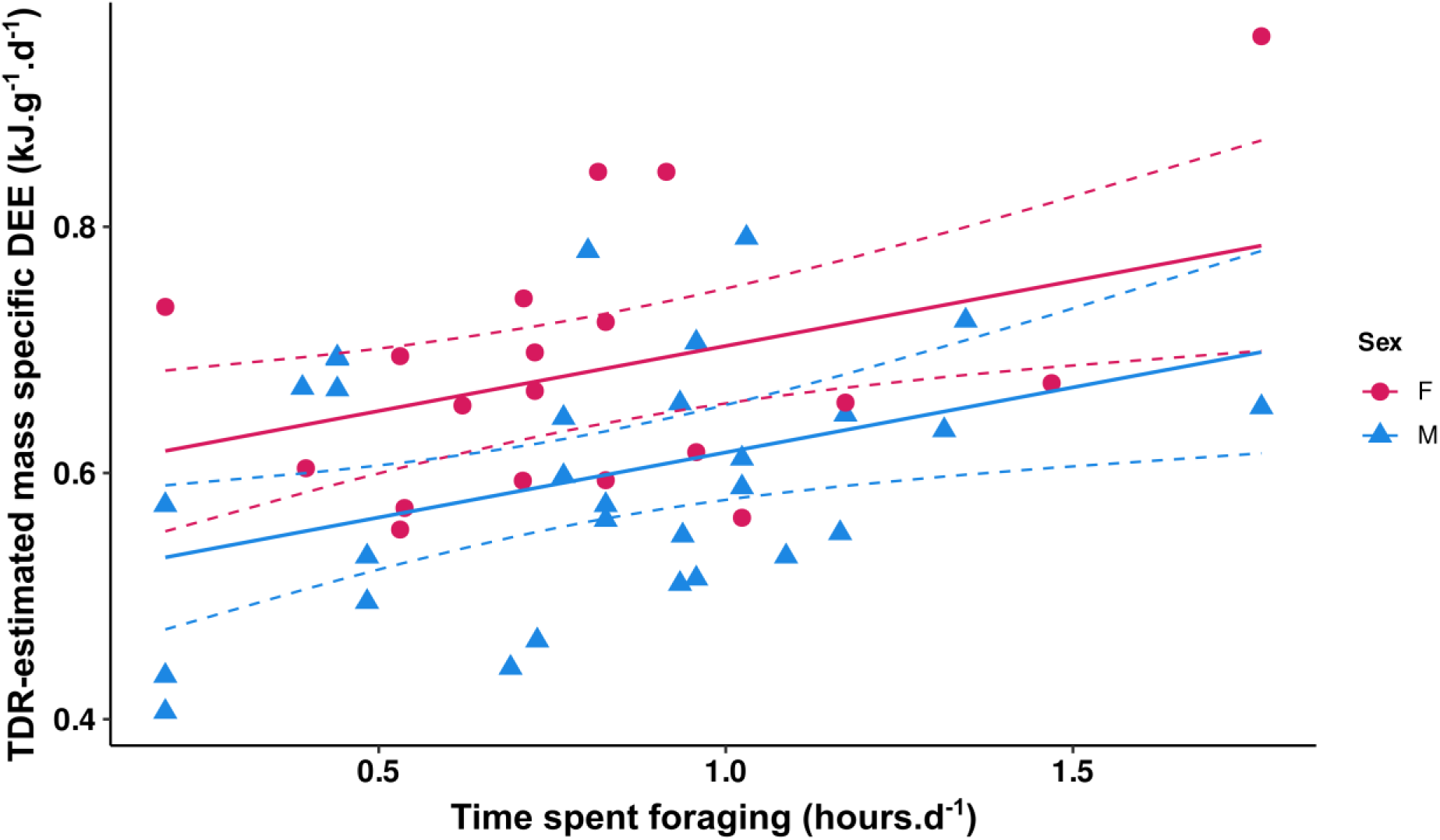
DEE increases with time spent foraging per day over the deployment. Dashed lines represent the 95% confidence interval around the regression.

## 4. Discussion

We present a novel method to quantify energetics (i.e energy acquisition and expenditure) based on depth data recorded from diving marine predators. We demonstrate the reliability of this method by using foraging trips from 48 Adélie penguins during the chick-rearing stage in the 2018-19 breeding-season.

Recordings from depth data provide information on animal movement in just one spatial dimension (y-axis), unlike accelerometry, which covers three spatial dimensions (x, y and z-axis). Furthermore, the temporal resolution of depth data is lower (1 Hz for TDR) compared to that of accelerometers (25 Hz or more). Yet, despite the lower spatial and temporal resolution, our results show that solely based on depth data, the machine learning algorithm RF can be used to identify fine-scale (1 Hz) foraging behaviour (*e.g* foraging events, prey encounters) with a good accuracy when trained from accelerometer data (accuracy = 0.83).

Our RF model was able to predict the foraging pattern detected with the accelerometry. However, the model grouped short prey catching attempts in single foraging bouts, which was expected given the lower resolution of depth data. Therefore, further investigation (e.g. using video cameras for example) is needed to assess what is being caught during these prey catching attempts (Del Caño et al., 2021; Sutton et al., 2020). Historical proxies like bottom duration or number of undulations were solely used to estimate proxies of foraging intensity without knowledge of when foraging was performed (Bost et al., 2007; Lescroël et al., 2021). Moreover, these historical proxies rely on the hypothesis that foraging is occurring mainly, if not totally, during the bottom phase of the dive (Deagle et al., 2008; Falk et al., 2000). Yet, accelerometry (Chimienti et al., 2022), oesophagus sensors (Ropert-Coudert et al., 2000) and video data (Del Caño et al., 2021; Sutton et al., 2020) have shown that penguins also feed during the ascent phase of their dive. By training our RF on foraging detected from accelerometry, our method was able to more precisely identify when feeding is happening using depth data, even during the ascent phase of dives, which was not possible with historical proxies. Moreover, the method presented in this paper presents the advantage of not needing arbitrary thresholds to define states such as bottom-phase or undulations, which could allow its application on other diving marine predators.

Overall, our method could likely be applied to other marine predators with TDR and accelerometer data availability to train species-specific RF algorithms. Indeed, the metrics used in this paper to estimate foraging are a simple and objective way of describing movement of diving predators. In addition to outperforming historical proxies like undulations and dive bottom phase duration, using this method would have the advantage to provide a simplified and unbiased comparison between studies. Yet, further investigations are needed to evaluate how our algorithm efficiently detects foraging for different types of prey, as foraging tactics and movement can change depending on the prey (Bowen et al., 2002). Krill is the main prey item of Adélie penguins, but they are also known to forage on fishes or squids (Ratcliffe and Trathan, 2012). Therefore, our dataset is likely to contain foraging of other prey than krill. Hence, our algorithm might also be set to investigate foraging of other penguin species if the metrics derived from TDR data are similar.

Moreover, we found that DEE increased with the proportion of time spent foraging per day. Because foraging is costly for diving marine predators (Jeanniard-du-Dot et al., 2017; Yeates et al., 2007), this correlation supports the idea that our methodology is reliable to estimate the energetics of Adélie penguins, and the benefits of studying energetics as both intake and expenditure rather than focusing on expenditure. Therefore, even if estimated time spent foraging needs further validation, our innovative methodology should improve our ability to study individual energetic trade-offs at large spatial and temporal scales. Here we identified a positive linear relationship between time spent foraging and DEE. Further studies could relate foraging duration to energy intake using mass gain measures and diet energy content estimation to better describe the extent of that relationship. Alternatively, validation of foraging detected through accelerometer using camera logger (Sutton et al., 2020; Watanabe et al., 2019) could link our estimated time spent foraging to energy intake.

With a R² of 0.70, our low-resolution TDR-based model provides comparable results to other DLW calibrations based on high-resolution accelerometers on marine and terrestrial species like Adélie penguin (R² = 0.75) (Hicks et al., 2020) or little penguins (R² = 0.78) (Sutton et al., 2021) or polar bear (R² = 0.70) (Pagano and Williams, 2019). This equation could be applied to the numerous Adélie penguin long-term TDR data collected across Antarctica (Cimino et al., 2023; Lescroël et al., 2023; Riaz et al., 2020), therefore offering a great opportunity to reliably investigate regional variations in energy budgets. In line with what was previously known (Hicks et al., 2020), we showed that assigning different calibration coefficients to different behaviour enhanced predictive power of our model. Our model shows penguins mostly expend their energy while diving and transiting (*i.e.* Time_Dive_ and Vertical movement_Sub-surface_ eqn.3). Also, despite being retained in the best model, sex was not a significant term and was not present in the second best performing model (delta AICc = 0.42), making our framework applicable in scenarios where individual sex is not known.

It is important to note that during the studied season, there was open water accessible next to the colony, and therefore, penguins did not have to walk long distances to access open water, reducing their energy expenditure (Watanabe et al., 2020). Therefore, our calibration might not reflect energy expenditure in areas and years when long walking periods are needed to access open water. Yet, the proposed model could still be relevant when focusing on the at-sea part of the foraging trips. Further investigations would be needed to assess if our model can also be used to estimate at-sea energy expenditure during the other phase of the breeding period, incubation, during which foraging trips are typically longer (10-15 days) and diving behaviour is different with shallower and less frequent dives (Chappell et al., 1993a; Lescroël et al., 2023). Yet, in their work Chappell *et al*. (Chappell et al., 1993b) found that the field metabolic rate of Adélie penguins was not significantly different in incubation and chick-rearing.

Historically, TDR had been the first type of logger deployed on wild animals (Kooyman, 1965). The described method can easily be applicable to other marine predators, with often longer time-series available for TDR data compared to accelerometers, allowing the study of long-term trends. With environmental changes affecting energy availability (Duncan et al., 2015), these long-term estimations of energetics are crucial to decipher how changing environmental conditions will affect individual life history strategies.

In conclusion, we show that lower-resolution TDR data can be used to estimate energetics similarly to accelerometers. Our results demonstrate how the application of machine learning approaches allows researchers to re-analyse datasets and more accurately predict energetics of diving marine predators compared to historical methods. Foraging activity is expected to be related to mass gain (Lescroël et al., 2021). Therefore, with prior knowledge of a species diet, calibrating predicted time spent foraging with mass gain could allow us to refine this simple framework by estimating energy intake directly. Because it’s based on the widely used TDR data, this framework could be applied to several long-term marine species monitoring programs. This would allow researchers to study how extrinsic (e.g. environmental variations) and intrinsic (e.g. body condition) variables impact individuals’ energetic trade-offs.

## Acknowledgements

We thank the Institut Polaire Francais Paul-Emile Victor (IPEV program P1091) for its financial and logistical support.

## Authors contribution

BD, AK, YRC and MC conceived the ideas and designed the methodology. OH, DW, CM, FA collected the data. BD analysed the data and wrote the manuscript. All authors contributed critically to the drafts and gave final approval for publication.

## Conflicts of interests statement

The authors declare no conflicts of interests

## Funding

This project has received funding from the European Union’s Horizon 2020 research and innovation programme under the Marie Sklodowska-Curie grant agreement No 890284, “Modelling Foraging Fitness in Marine predators (MuFFIN)”, awarded to Marianna Chimienti, WWF-UK and PEW foundation. This work is part of a PhD project funded by a grant from the French ministry of higher education and research awarded to Benjamin Dupuis.

## Data availability

All code used in this paper are available at https://github.com/bendps/tdr2nrj

## Diversity and inclusion statement

All authors were engaged early on with the research and study design to ensure that the diverse sets of perspectives they represent was considered from the onset. Whenever relevant, literature published by scientists from different regions was cited.

## Notes

https://github.com/bendps/tdr2nrj

### Competing Interest Statement

The authors have declared no competing interest.

## References

1. Ainley, D., 2002. The Adélie Penguin: Bellwether of Climate Change, in: The Adélie Penguin. Columbia University Press. 10.7312/ainl12306

2. Allegue, H., Réale, D., Picard, B., Guinet, C., 2023. Track and dive-based movement metrics do not predict the number of prey encountered by a marine predator. Movement Ecology 11, 3. 10.1186/s40462-022-00361-2

3. Barbraud, C., Delord, K., Bost, C.A., Chaigne, A., Marteau, C., Weimerskirch, H., 2020. Population trends of penguins in the French Southern Territories. Polar Biol 43, 835–850. 10.1007/s00300-020-02691-6

4. Bestley, S., Jonsen, I.D., Hindell, M.A., Harcourt, R.G., Gales, N.J., 2015. Taking animal tracking to new depths: synthesizing horizontal–vertical movement relationships for four marine predators. Ecology 96, 417–427. 10.1890/14-0469.1

5. Bost, C.A., Handrich, Y., Butler, P.J., Fahlman, A., Halsey, L.G., Woakes, A.J., Ropert-Coudert, Y., 2007. Changes in dive profiles as an indicator of feeding success in king and Adélie penguins. Deep Sea Research Part II: Topical Studies in Oceanography 54, 248–255. 10.1016/j.dsr2.2006.11.007

6. Bowen, W.D., Tully, D., Boness, D.J., Bulheier, B.M., Marshall, G.J., 2002. Prey-dependent foraging tactics and prey profitability in a marine mammal. Marine Ecology Progress Series 244, 235–245. 10.3354/meps244235

7. Brisson-Curadeau, É., Handrich, Y., Elliott, K.H., Bost, C.-A., 2021. Accelerometry predicts prey-capture rates in the deep-diving king penguin Aptenodytes patagonicus. Mar Biol 168, 156. 10.1007/s00227-021-03968-y

8. Brown, J.H., Gillooly, J.F., Allen, A.P., Savage, V.M., West, G.B., 2004. Toward a Metabolic Theory of Ecology. Ecology 85, 1771–1789. 10.1890/03-9000

9. Burger, J.R., Hou, C., Brown, J.H., 2019. Toward a metabolic theory of life history. Proceedings of the National Academy of Sciences 116, 26653–26661. 10.1073/pnas.1907702116

10. Byrne, B., Kort, S.R. de, Pedley, S.M., 2022. Leafcutter ants adjust foraging behaviours when exposed to noise disturbance. PLOS ONE 17, e0269517. 10.1371/journal.pone.0269517

11. Carter, M.I.D., Russell, D.J.F., Embling, C.B., Blight, C.J., Thompson, D., Hosegood, P.J., Bennett, K.A., 2017. Intrinsic and extrinsic factors drive ontogeny of early-life at-sea behaviour in a marine top predator. Sci Rep 7, 15505. 10.1038/s41598-017-15859-8

12. Chappell, M.A., Shoemaker, V.H., Janes, D.N., Bucher, T.L., Maloney, S.K., 1993a. Diving Behavior During Foraging in Breeding Adelie Penguins. Ecology 74, 1204–1215. 10.2307/1940491

13. Chappell, M.A., Shoemaker, V.H., Janes, D.N., Maloney, S.K., Bucher, T.L., 1993b. Energetics of Foraging in Breeding Adelie Penguins. Ecology 74, 2450–2461. 10.2307/1939596

14. Chevallay, M., Guinet, C., Jeanniard-Du-Dot, T., 2022. Should I stay or should I go? Behavioral adjustments of fur seals related to foraging success. Behavioral Ecology 33, 634–643. 10.1093/beheco/arac012

15. Chimienti, M., Cornulier, T., Owen, E., Bolton, M., Davies, I.M., Travis, J.M.J., Scott, B.E., 2016. The use of an unsupervised learning approach for characterizing latent behaviors in accelerometer data. Ecology and Evolution 6, 727–741. 10.1002/ece3.1914

16. Chimienti, M., Kato, A., Hicks, O., Angelier, F., Beaulieu, M., Ouled-Cheikh, J., Marciau, C., Raclot, T., Tucker, M., Wisniewska, D.M., Chiaradia, A., Ropert-Coudert, Y., 2022. The role of individual variability on the predictive performance of machine learning applied to large bio-logging datasets. Sci Rep 12, 19737. 10.1038/s41598-022-22258-1

17. Chivers, L.S., Lundy, M.G., Colhoun, K., Newton, S.F., Houghton, J.D.R., Reid, N., 2012. Foraging trip time-activity budgets and reproductive success in the black-legged kittiwake. Marine Ecology Progress Series 456, 269–277. 10.3354/meps09691

18. Cimino, M.A., Conroy, J.A., Connors, E., Bowman, J., Corso, A., Ducklow, H., Fraser, W., Friedlaender, A., Kim, H.H., Larsen, G.D., Moffat, C., Nichols, R., Pallin, L., Patterson-Fraser, D., Roberts, D., Roberts, M., Steinberg, D.K., Thibodeau, P., Trinh, R., Schofield, O., Stammerjohn, S., 2023. Long-term patterns in ecosystem phenology near Palmer Station, Antarctica, from the perspective of the Adélie penguin. Ecosphere 14, e4417. 10.1002/ecs2.4417

19. Deagle, B.E., Gales, N.J., Hindell, M.A., 2008. Variability in foraging behaviour of chick-rearing macaroni penguins Eudyptes chrysolophus and its relation to diet. Marine Ecology Progress Series 359, 295–309. 10.3354/meps07307

20. Del Caño, M., Quintana, F., Yoda, K., Dell’Omo, G., Blanco, G.S., Gómez-Laich, A., 2021. Fine-scale body and head movements allow to determine prey capture events in the Magellanic Penguin (Spheniscus magellanicus). Mar Biol 168, 84. 10.1007/s00227-021-03892-1

21. Duncan, C., Chauvenet, A.L.M., Brown, M.E., Pettorelli, N., 2015. Energy availability, spatio-temporal variability and implications for animal ecology. Diversity Distrib. 21, 290–301. 10.1111/ddi.12270

22. Egert-Berg, K., Handel, M., Goldshtein, A., Eitan, O., Borissov, I., Yovel, Y., 2021. Fruit bats adjust their foraging strategies to urban environments to diversify their diet. BMC Biol 19, 1–11. 10.1186/s12915-021-01060-x

23. English, H.M., Börger, L., Kane, A., Ciuti, S., 2024. Advances in biologging can identify nuanced energetic costs and gains in predators. Mov Ecol 12, 1–17. 10.1186/s40462-024-00448-y

24. Falk, K., Benvenuti, S., Dall’antonia, L., Kampp, K., Ribolini, A., 2000. Time allocation and foraging behaviour of chick-rearing Brünnich’s Guillemots Uria lomvia in high-arctic Greenland. Ibis 142, 82–92. 10.1111/j.1474-919X.2000.tb07687.x

25. Forcada, J., Trathan, P.N., 2009. Penguin responses to climate change in the Southern Ocean. Global Change Biology 15, 1618–1630. 10.1111/j.1365-2486.2009.01909.x

26. Froget, G., Butler, P.J., Woakes, A.J., Fahlman, A., Kuntz, G., Le Maho, Y., Handrich, Y., 2004. Heart rate and energetics of free-ranging king penguins (Aptenodytes patagonicus). J Exp Biol 207, 3917–3926. 10.1242/jeb.01232

27. Gallon, S., Bailleul, F., Charrassin, J.-B., Guinet, C., Bost, C.-A., Handrich, Y., Hindell, M., 2013. Identifying foraging events in deep diving southern elephant seals, *Mirounga leonina*, using acceleration data loggers. Deep Sea Research Part II: Topical Studies in Oceanography, Fourth International Symposium on Bio-logging Science 88–89, 14–22. 10.1016/j.dsr2.2012.09.002

28. Gower, C.N., Garrott, R.A., White, P.J., Watson, F.G.R., Cornish, S.S., Becker, M.S., 2008. Chapter 18 Spatial Responses of Elk to Wolf Predation Risk: Using the Landscape to Balance Multiple Demands, in: Garrott, R.A., White, P.J., Watson, F.G.R. (Eds.), Terrestrial Ecology, The Ecology of Large Mammals in Central Yellowstone. Elsevier, pp. 373–399. 10.1016/S1936-7961(08)00218-2

29. Hazen, E.L., Abrahms, B., Brodie, S., Carroll, G., Jacox, M.G., Savoca, M.S., Scales, K.L., Sydeman, W.J., Bograd, S.J., 2019. Marine top predators as climate and ecosystem sentinels. Frontiers in Ecology and the Environment 17, 565–574. 10.1002/fee.2125

30. Hicks, O., Kato, A., Angelier, F., Wisniewska, D.M., Hambly, C., Speakman, J.R., Marciau, C., Ropert-Coudert, Y., 2020. Acceleration predicts energy expenditure in a fat, flightless, diving bird. Sci Rep 10, 21493. 10.1038/s41598-020-78025-7

31. Jeanniard-du-Dot, T., Guinet, C., Arnould, J.P.Y., Speakman, J.R., Trites, A.W., 2017. Accelerometers can measure total and activity-specific energy expenditures in free-ranging marine mammals only if linked to time-activity budgets. Functional Ecology 31, 377–386. 10.1111/1365-2435.12729

32. Kokubun, N., Emmerson, L., McInnes, J., Wienecke, B., Southwell, C., 2021. Sea-ice and density-dependent factors affecting foraging habitat and behaviour of Adélie penguins throughout the breeding season. Mar Biol 168, 97. 10.1007/s00227-021-03899-8

33. Kokubun, N., Kim, J.-H., Shin, H.-C., Naito, Y., Takahashi, A., 2011. Penguin head movement detected using small accelerometers: a proxy of prey encounter rate. Journal of Experimental Biology 214, 3760–3767. 10.1242/jeb.058263

34. Kooyman, G.L., 1965. Techniques used in measuring diving capacities of Weddell Seals. Polar Record 12, 391–394. 10.1017/S003224740005484X

35. Kressler, M.M., Dall, S.R.X., Sherley, R.B., 2023. A framework for studying ecological energy in the contemporary marine environment. ICES Journal of Marine Science fsad082. 10.1093/icesjms/fsad082

36. Kuhn, M., Wickham, H., 2020. Tidymodels: a collection of packages for modeling and machine learning using tidyverse principles.

37. Kursa, M.B., Rudnicki, W.R., 2010. Feature Selection with the Boruta Package. Journal of Statistical Software 36, 1–13.

38. Leroy, C.C., Parthiot, F., 1998. Depth-pressure relationships in the oceans and seas. The Journal of the Acoustical Society of America 103, 1346–1352. 10.1121/1.421275

39. Lescroël, A., Schmidt, A., Ainley, D.G., Dugger, K.M., Elrod, M., Jongsomjit, D., Morandini, V., Winquist, S., Ballard, G., 2023. High-resolution recording of foraging behaviour over multiple annual cycles shows decline in old Adélie penguins’ performance. Proceedings of the Royal Society B: Biological Sciences 290, 20222480. 10.1098/rspb.2022.2480

40. Lescroël, A., Schmidt, A., Elrod, M., Ainley, D.G., Ballard, G., 2021. Foraging dive frequency predicts body mass gain in the Adélie penguin. Sci Rep 11, 22883. 10.1038/s41598-021-02451-4

41. Lüdecke, D., 2023. sjPlot: Data Visualization for Statistics in Social Science.

42. Machado-Gaye, A.L., Kato, A., Chimienti, M., Gobel, N., Ropert-Coudert, Y., Barbosa, A., Soutullo, A., 2024. Using latent behavior analysis to identify key foraging areas for Adélie penguins in a declining colony in West Antarctic Peninsula. Mar Biol 171, 69. 10.1007/s00227-024-04390-w

43. Marciau, C., Raclot, T., Bestley, S., Barbraud, C., Delord, K., Hindell, M.A., Kato, A., Parenteau, C., Poupart, T., Ribout, C., Ropert-Coudert, Y., Angelier, F., 2023. Body condition and corticosterone stress response, as markers to investigate effects of human activities on Adélie penguins (Pygoscelis adeliae). Front. Ecol. Evol. 11. 10.3389/fevo.2023.1099028

44. Michelot, C., Kato, A., Raclot, T., Shiomi, K., Goulet, P., Bustamante, P., Ropert-Coudert, Y., 2020. Sea-ice edge is more important than closer open water access for foraging Adélie penguins: evidence from two colonies. Mar. Ecol. Prog. Ser. 640, 215–230. 10.3354/meps13289

45. Mogensen, S., Post, J.R., 2012. Energy allocation strategy modifies growth–survival trade-offs in juvenile fish across ecological and environmental gradients. Oecologia 168, 923–933. 10.1007/s00442-011-2164-0

46. Morano, S., Stewart, K.M., Sedinger, J.S., Nicolai, C.A., Vavra, M., 2013. Life-history strategies of North American elk: trade-offs associated with reproduction and survival. Journal of Mammalogy 94, 162–172. 10.1644/12-MAMM-A-074.1

47. Nagy, K.A., Girard, I.A., Brown, T.K., 1999. ENERGETICS OF FREE-RANGING MAMMALS, REPTILES, AND BIRDS. Annual Review of Nutrition 19, 247–277. 10.1146/annurev.nutr.19.1.247

48. Noakes, M.J., Smit, B., Wolf, B.O., McKechnie, A.E., 2013. Thermoregulation in African Green Pigeons (Treron calvus) and a re-analysis of insular effects on basal metabolic rate and heterothermy in columbid birds. J Comp Physiol B 183, 969–982. 10.1007/s00360-013-0763-2

49. Pagano, A.M., Williams, T.M., 2019. Estimating the energy expenditure of free-ranging polar bears using tri-axial accelerometers: A validation with doubly labeled water. Ecology and Evolution 9, 4210–4219. 10.1002/ece3.5053

50. Pichler, M., Hartig, F., 2023. Machine learning and deep learning—A review for ecologists. Methods in Ecology and Evolution 14, 994–1016. 10.1111/2041-210X.14061

51. Pontzer, H., McGrosky, A., 2022. Balancing growth, reproduction, maintenance, and activity in evolved energy economies. Current Biology 32, R709–R719. 10.1016/j.cub.2022.05.018

52. R Core Team, 2023. R: A Language and Environment for Statistical Computing. R Foundation for Statistical Computing, Vienna, Austria.

53. Ratcliffe, N., Trathan, P., 2012. A review of the diet and at-sea distribution of penguins breeding within the CAMLR Convention Area. CCAMLR Science 19, 75–114.

54. Riaz, J., Bestley, S., Wotherspoon, S., Freyer, J., Emmerson, L., 2020. From trips to bouts to dives: temporal patterns in the diving behaviour of chick-rearing Adélie penguins, East Antarctica. Marine Ecology Progress Series 654, 177–194. 10.3354/meps13519

55. Ropert-Coudert, Y., Baudat, J., Kurita, M., Bost, C.-A., Kato, A., Le Maho, Y., Naito, Y., 2000. Validation of oesophagus temperature recording for detection of prey ingestion on captive Adélie penguins (Pygoscelis adeliae). Marine Biology 137, 1105–1110. 10.1007/s002270000414

56. Ropert-Coudert, Y., Beaulieu, M., Hanuise, N., Kato, A., 2009. Diving into the world of biologging. Endangered Species Research 10, 21–27. 10.3354/esr00188

57. Ropert-Coudert, Y., Kato, A., Baudat, J., Bost, C.-A., Le Maho, Y., Naito, Y., 2001. Feeding strategies of free-ranging Adélie penguins Pygoscelis adeliae analysed by multiple data recording. Polar Biology 24, 460–466. 10.1007/s003000100234

58. Ropert-Coudert, Y., Wilson, R.P., Yoda, K., Kato, A., 2007. Assessing performance constraints in penguins with externally-attached devices. Marine Ecology Progress Series 333, 281–289. 10.3354/meps333281

59. Sergio, F., Caro, T., Brown, D., Clucas, B., Hunter, J., Ketchum, J., McHugh, K., Hiraldo, F., 2008. Top Predators as Conservation Tools: Ecological Rationale, Assumptions, and Efficacy. Annu. Rev. Ecol. Evol. Syst. 39, 1–19. 10.1146/annurev.ecolsys.39.110707.173545

60. Steinhart, G.B., Sandrene, M.E., Weaver, S., Stein, R.A., Marschall, E.A., 2005. Increased parental care cost for nest-guarding fish in a lake with hyperabundant nest predators. Behavioral Ecology 16, 427–434. 10.1093/beheco/ari006

61. Sutton, G., Pichegru, L., Botha, J.A., Kouzani, A.Z., Adams, S., Bost, C.A., Arnould, J.P.Y., 2020. Multi-predator assemblages, dive type, bathymetry and sex influence foraging success and efficiency in African penguins. PeerJ 8, e9380. 10.7717/peerj.9380

62. Sutton, G.J., Botha, J.A., Speakman, J.R., Arnould, J.P.Y., 2021. Validating accelerometry-derived proxies of energy expenditure using the doubly labelled water method in the smallest penguin species. Biol Open 10, bio055475. 10.1242/bio.055475

63. Sydeman, W.J., Poloczanska, E., Reed, T.E., Thompson, S.A., 2015. Climate change and marine vertebrates. Science 350, 772–777. 10.1126/science.aac9874

64. Viviant, M., Monestiez, P., Guinet, C., 2014. Can We Predict Foraging Success in a Marine Predator from Dive Patterns Only? Validation with Prey Capture Attempt Data. PLOS ONE 9, e88503. 10.1371/journal.pone.0088503

65. Watanabe, Y.Y., Ito, K., Kokubun, N., Takahashi, A., 2020. Foraging behavior links sea ice to breeding success in Antarctic penguins. Science Advances 6, eaba4828. 10.1126/sciadv.aba4828

66. Watanabe, Y.Y., Payne, N.L., Semmens, J.M., Fox, A., Huveneers, C., 2019. Hunting behaviour of white sharks recorded by animal-borne accelerometers and cameras. Marine Ecology Progress Series 621, 221–227. 10.3354/meps12981

67. Wilson, R.P., Börger, L., Holton, M.D., Scantlebury, D.M., Gómez-Laich, A., Quintana, F., Rosell, F., Graf, P.M., Williams, H., Gunner, R., Hopkins, L., Marks, N., Geraldi, N.R., Duarte, C.M., Scott, R., Strano, M.S., Robotka, H., Eizaguirre, C., Fahlman, A., Shepard, E.L.C., 2020. Estimates for energy expenditure in free-living animals using acceleration proxies: A reappraisal. Journal of Animal Ecology 89, 161–172. 10.1111/1365-2656.13040

68. Wright, M.N., Ziegler, A., 2017. ranger: A Fast Implementation of Random Forests for High Dimensional Data in C++ and R. Journal of Statistical Software 77, 1–17. 10.18637/jss.v077.i01

69. Yeates, L.C., Williams, T.M., Fink, T.L., 2007. Diving and foraging energetics of the smallest marine mammal, the sea otter (Enhydra lutris). Journal of Experimental Biology 210, 1960–1970. 10.1242/jeb.02767

