## Supplementaries for "Innovative Use of Depth Data to Estimate Energy Intake and Expenditure in Adélie Penguins"

### Supplementary information

**Figure S1.** Output from the most parsimonious model to predict DLW-derived DEE. A. Daily amount of vertical movement in the subsurface phase. B. Daily number of hours spent diving. Dashed lines represent the 95% confidence interval around the regression. Dots represent the partial residuals of the model.

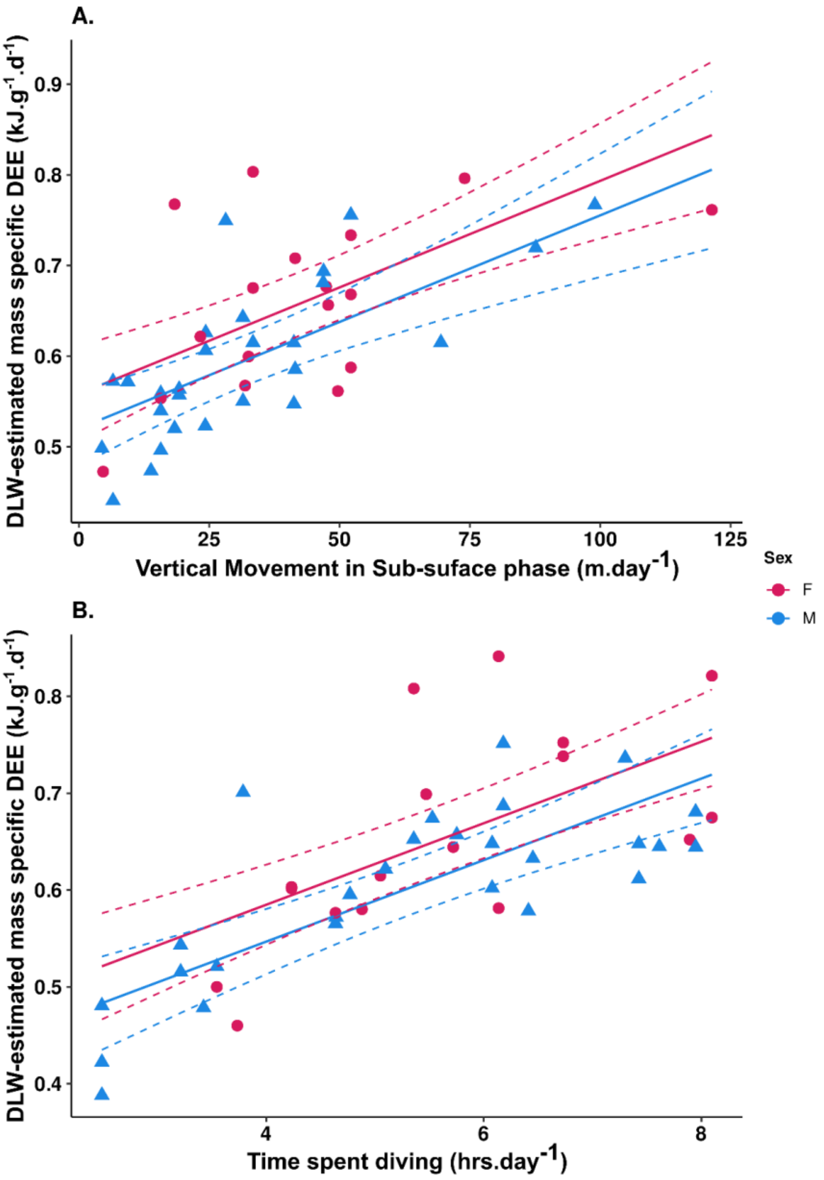

**Figure S2.** Relative variable importance from the random forest predicting accelerometry-based time spent foraging from depth-derived parameters.

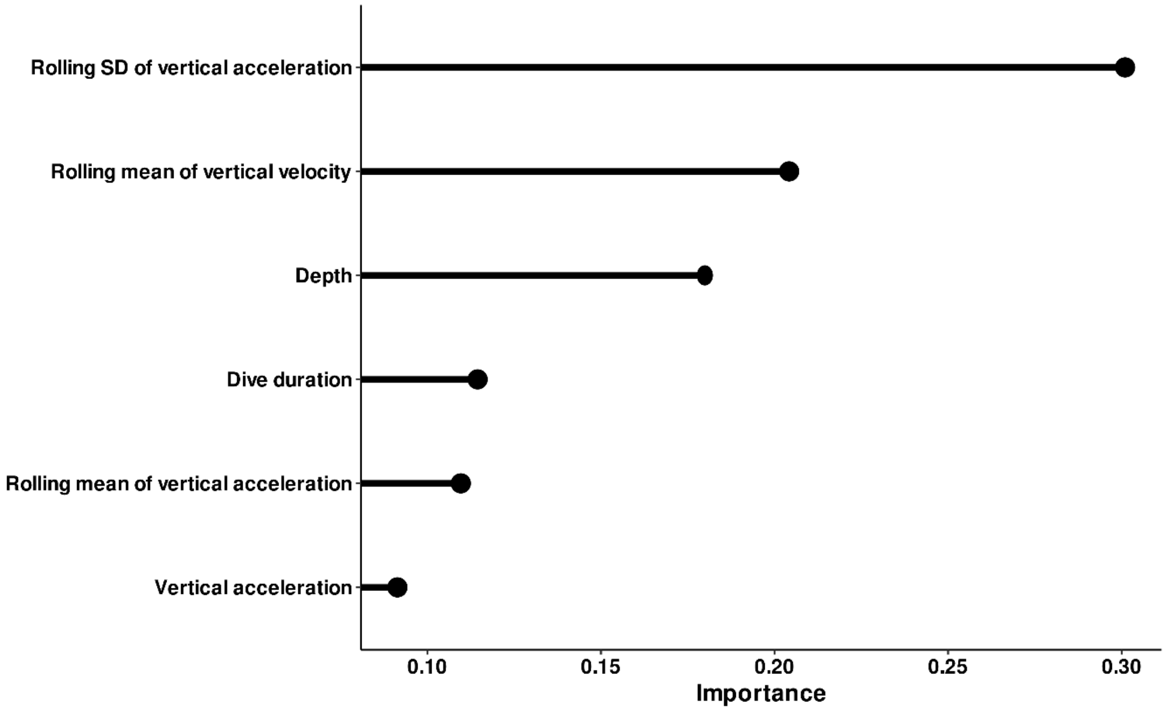

**Table S1.** Most parsimonious model summary for DLW-derived DEE.

| **Characteristic** | **Beta** | **95% CI**[1] | **p-value** | **VIF**[1] |
|---|---|---|---|---|
| *(Intercept)* | 0.33 | 0.25, 0.41 | <0.001 | |
| *Time(dive)* | 0.04 | 0.03, 0.06 | <0.001 | 1.1 |
| *Vertical movement(sub-surface)* | 0.00 | 0.00, 0.00 | <0.001 | 1.1 |
| *Sex* | | | | 1.1 |
| F | — | — | | |
| M | -0.04 | -0.08, 0.01 | 0.10 | |

[1]CI = Confidence Interval, VIF = Variance Inflation Factor

**Table S2.** Most parsimonious model summary for predicting time spent foraging.

| Characteristic | Beta | 95% CI[1] | p-value |
|---|---|---|---|
| *(Intercept)* | 0.38 | 0.044, 0.72 | 0.028 |
| *Time(TDR-estimated hunting)* | 0.78 | 0.62, 0.94 | <0.001 |

[1]CI = Confidence Interval

**Table S3.** Model selection and coefficients of the models estimating relationship between time spent foraging and DEE.

| (Intercept) | Time spent foraging per day | Sex | *df* | logLik | AICc | ΔAICc | weight |
|---|---|---|---|---|---|---|---|
| 0.60 | 0.11 | + | 4 | 46.69 | -84.48 | 0.00 | 0.90 |
| 0.68 | | + | 3 | 42.89 | -79.25 | 5.23 | 0.07 |
| 0.54 | 0.11 | | 3 | 42.15 | -77.76 | 6.72 | 0.03 |
| 0.63 | | | 2 | 38.81 | -73.35 | 11.13 | 0.00 |
